# Comparing the speed of action initiation and action inhibition

**DOI:** 10.1101/2022.06.29.497798

**Authors:** Yue Du, Alexander D. Forrence, Delaney M. Metcalf, Adrian M. Haith

## Abstract

Much work has examined the process of canceling or inhibiting an impending action. However, the converse process of initiating a movement has, comparatively, been much less studied. Action initiation and action inhibition are generally considered to be independent and qualitatively distinct processes, while the exact relationship between these processes remains unclear. One respect in which action initiation and action inhibition are often thought to differ is in their speed; action inhibition is typically considered to be faster than action initiation, which allows for impending actions to be inhibited before they are initiated. This apparent contrast is, however, largely observed in tasks in which there is much greater urgency to inhibit an action that there is to initiate an action. This asymmetry in the urgency between action initiation and action inhibition, as well as other asymmetries in how action initiation and inhibition are cued, make it impossible to compare their relative time courses. Here, we demonstrate that, when action initiation and action inhibition are measured under conditions that are matched as closely as possible, their speed is the same. In light of this, we suggest that action initiation and action inhibition may not necessarily be qualitatively distinct processes, but may instead reflect two opposing states of a single process supporting a decision about whether to act or not. This perspective carries significant implications for computational models and presumed neural mechanisms of action control.

## Introduction

Initiating a movement quickly when needed is a fundamental aspect of action control, but its basic mechanisms and properties are not completely understood. In particular, action initiation has not been examined nearly as much as action inhibition – the ability to cancel or inhibit an impending voluntary action moments before it is initiated (Logan and Cowan, 1984). Decades of research on action inhibition has led to a detailed understanding of its neural basis, its role in executive function, and the nature of its impairment in psychopathology and neuropathology (Aron et al., 2014; Dunovan et al., 2015; Hannah and Aron, 2021; Oosterlaan et al., 1998; Verbruggen and Logan, 2008; Wiecki and Frank, 2013). We currently lack a similarly comprehensive understanding of the properties and mechanisms of generating an action, and little sense of exactly how it might relate to action inhibition.

The process of inhibiting an impending action has mostly been studied using the stop-signal task. In this task, people are asked to respond to an imperative “go” stimulus as quickly as possible with a button press (or other discrete or continuous actions) but must cancel this response in the event of a “stop” signal (Logan and Cowan, 1984; Verbruggen et al., 2019). Human behavior in such tasks is commonly interpreted through the independent race model in which a go process and a stop process accumulate independently towards a threshold. Whichever process first reaches the threshold determines whether or not the action will be generated (Logan et al., 2014; Logan and Cowan, 1984). In these models, the stop process generally accumulates much faster than the go process, accounting for the fact that actions can often successfully be stopped even when the stop signal appears much later than the go signal. This is consistent with experimental observations that it takes a shorter time to inhibit a response than to initiate one (Leunissen et al., 2017; Logan and Cowan, 1984; Slater-Hammel, 1960).

This apparent speed advantage of action inhibition over action initiation may not, however, reflect a general difference in their properties. Because studies using the stop-signal task primarily focus on action inhibition, they are often deliberately designed to slow responses to the go stimulus in order to promote successful stopping (e.g. by imposing a response choice or introducing stop/go cues through different sensory modalities) (e.g., Bissett and Logan, 2014; Matzke et al., 2018; Raud et al., 2020; Verbruggen et al., 2019; Verbruggen and Logan, 2017; Wadsley et al., 2022; Xu et al., 2015). Even when the task is not overtly designed to slow action initiation, however, there is still a significant asymmetry in the urgency to generate a response to the go cue and the urgency to cancel a response following the stop cue; cancellation of a response has to occur before the movement is initiated but, in contrast, a response can be initiated at any time following the presentation of the go cue. In fact, it can be advantageous to deliberately slow action initiation in order to accommodate the stopping behavior (Lappin and Eriksen, 1966; Leotti and Wager, 2010; Verbruggen and Logan, 2008). This asymmetry in urgency, which is inherent in the design of the stop-signal task, makes it impossible to directly compare the properties of action initiation and action inhibition within conventional stop signal tasks.

To more fairly compare the relative speed of action initiation and action inhibition, we performed an experiment to measure them in two separate tasks in which participants could generate an action or cancel an intended action. By measuring action initiation and inhibition in separate tasks, we were able to match the urgency as closely as possible. Key to our experimental design was to adopt a “timed-response” approach in which participants were trained to always respond at a prescribed time in each trial (Haith et al., 2016). By occasionally and unexpectedly switching the required behavior, either from requiring no response to requiring a response, or from requiring a response at the prescribed time to requiring no response (Coxon et al., 2006; Leunissen et al., 2017; Slater-Hammel, 1960), we were able to measure the time course of initiation and inhibition in symmetric conditions, allowing us to directly compare them.

## Results

Participants performed two tasks designed to assess how quickly they could generate or cancel a movement in response to an unexpected cue. Participants viewed a circle moving vertically downward to cross a horizonal line (Fig. 1) and were instructed to either press a button when the circle overlapped the target line, or do nothing, depending on the color of the circle as it crossed the line. By varying the initial color of the circle and unexpectedly changing the color during the trial (switch trials; 30% of all trials), we created two separate conditions to assess the speed of action initiation and action inhibition, respectively: In the Response-to-No-Response condition (R-to-NR), participants needed to rapidly abort an initially prepared response (similar to conventional stopping paradigms (Coxon et al., 2006; Slater-Hammel, 1960; Verbruggen et al., 2019)); in the No-Response-to-Response condition (NR-to-R) participants needed to rapidly initiate a response that they had not initially intended to.

**Fig. 1.**
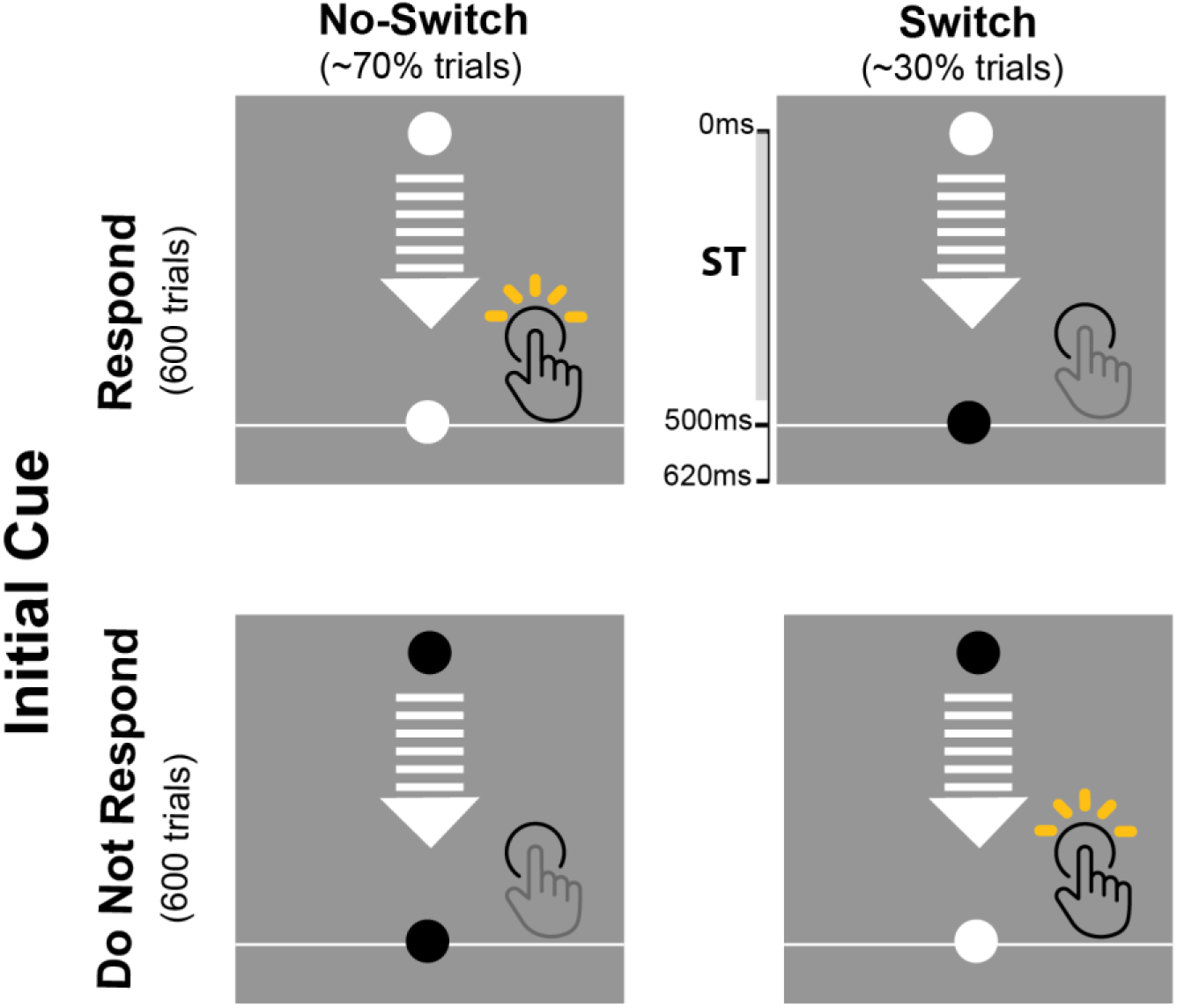
Experiment procedure. Participants were asked to either press a key or do nothing when a moving circle reached the target line. Whether or not a response was required in a given trial depended on the color of the circle (e.g., white = respond; black = do not respond). The actual circle color was counterbalanced across participants, controlling for the potential perceptual differences of black and white colors. The circle always started with the same color within each block of 100 trials. In ~70% of trials, the circle remained the same color throughout. However, in another ~30% of trials, the circle changed color before it hit the target line, forcing participants to cancel a preplanned response or to initiate a response when the circle crossed the line. By manipulating the time at which the circle color changed in each condition, we were able to compare people’s ability to stop themselves from generating a planned response (“R-to-NR” condition; upper panel) to the ability to rapidly generate a response (“NR-to-R” condition; lower panel). Participants completed 12 blocks of 100 trials, generating 204 switch trials in each condition.

The performance of one exemplar participant is shown in Fig. 2B. In switch trials of the R-to-NR condition, participants had to abort an impending response. The amount of time available to participants to inhibit their response, which we term the *allowed RT* (Figure 2A; Methods), depended on when the circle changed color, which varied from 50 ms to 500 ms before crossing the target line. When the allowed RT was very short (< 100ms), this participant almost always failed to abort their prepared response. However, at longer allowed RTs (> 300ms), this participant was able to correctly cancel the response in almost all trials. At intermediate allowed RTs, the participant was sometimes successful and sometimes unsuccessful in canceling their response. This behavior was broadly consistent with previous findings in stop-signal task (Coxon et al., 2006; Dunovan et al., 2015; Leunissen et al., 2017; Slater-Hammel, 1960; Verbruggen et al., 2019).

**Fig. 2.**
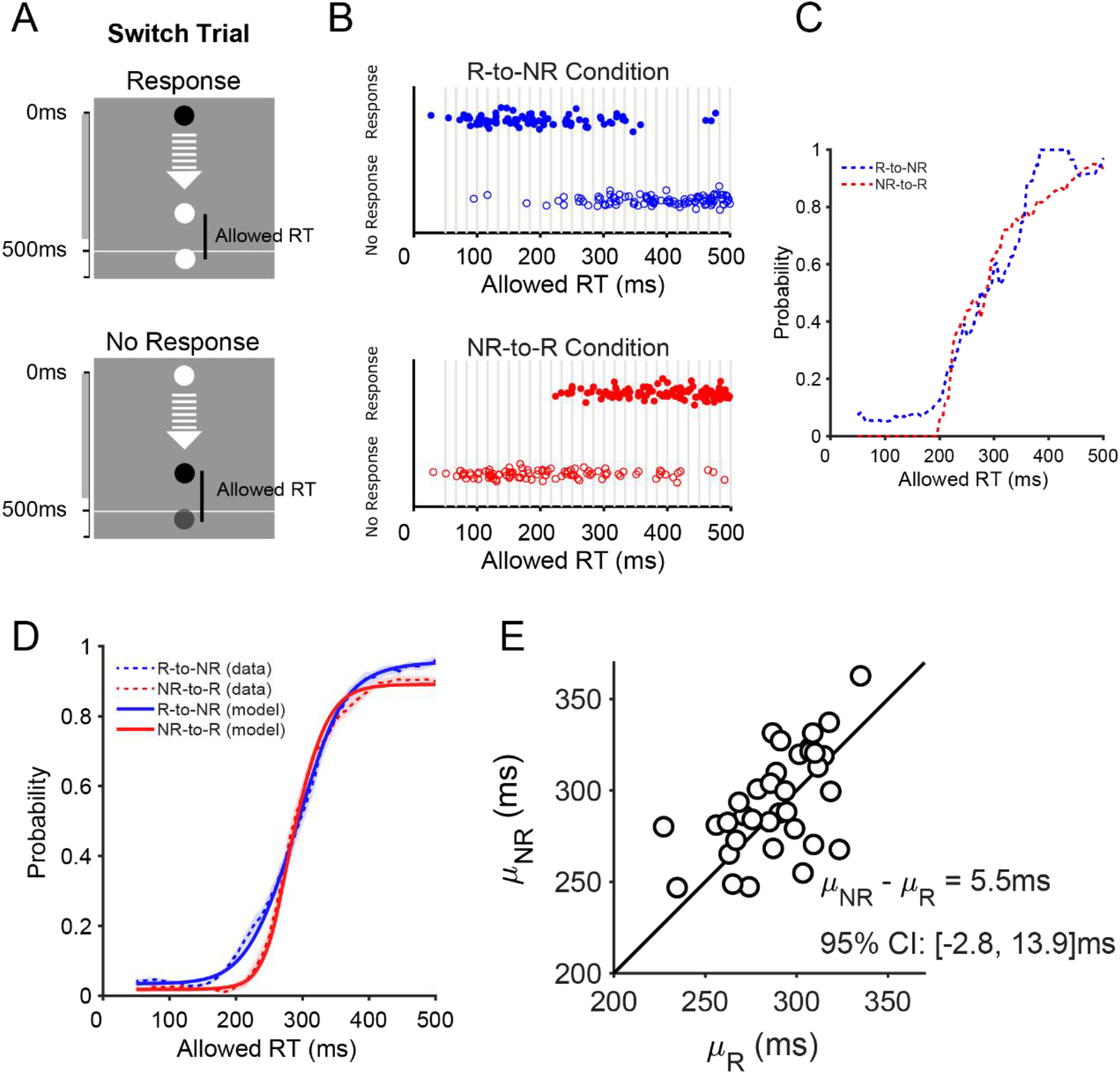
Cancelling an impending response is not faster than initiating one. A) The allowed RT was quantified as the time elapsed from color change to the time of the button press when participants generated a response. When no response occurred, the allowed RT was approximated as the time interval between the color change to the typical time of button presses in comparable trials (see Methods for more details). B) Behavior of one exemplar participant. In trials in which only a very short allowed RT was allowed, this participant consistently made the wrong choice as to whether to respond or not. When a longer allowed RT was allowed, this participant was able to consistently make the correct choice to respond or not. Vertical jitter was added to allow individual data points to be seen more easily. C) The raw data were used to construct speed–accuracy trade-offs, showing the probability of a correct choice as a function of allowed RT. D) Mean speed–accuracy trade-offs for each condition across all participants (dashed lines) were well captured by a computational model (solid lines) in which we assumed that the decision to respond or not could be thought of as a discrete event occurring at a random time 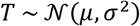 after the circle changes color. E) We used this model to estimate the average time needed to cancel an intended response *μ_NR_* in R-to-NR condition and the average time to initiate a response *μ_R_* in the NR-to-R condition. Across all participants, *μ_NR_* (294.5 ms ± 28.3 ms; mean ± s.d.) was not significantly different from *μ_R_* (289.0 ms ± 24.4 ms; mean ± s.d.).

Similar but complementary behavior was observed in the NR-to-R switch trials. This participant failed to initiate a timely response if the circle changed color shortly before it crossed the line. When allowed a longer RT to react to the color change, however, they always correctly generated a response as the circle crossed the target line.

From this raw response data, we constructed a speed–accuracy trade-off for each condition (R-to-NR and NR-to-R), based on a 50 ms sliding window on the allowed RT. This trade-off function describes the probability of correctly aborting a response (R-to-NR) or the probability of generating a successful response (NR-to-R) as a function of allowed RT (Fig. 2C). For the exemplar participant in Fig. 2B, the centers of the two speed–accuracy trade-off functions were both located at around 280 ms, indicating that the average times required to cancel a response (R-to-NR condition) or initiate a response (NR-to-R) were comparable. Averaged behavior across all participants (n = 35 out of 36; Methods; Figs. S1 and S2) showed the same pattern (Fig. 2D; Dashed lines), with comparable RTs required to initiate a response and to cancel the initiation of a response.

To more precisely quantify how quickly participants could cancel a response in the R-to-NR condition, we considered a simple model – similar to the classic race model of stopping behavior (Logan and Cowan, 1984) – in which we assumed that the cancellation of a response was a discrete event occurring at a random time 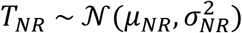 after the circle changed color. On a given trial, if the time needed to cancel a response (*T_NR_*) was shorter than the allowed RT, participants would successfully avoid generating a response. But, if the required time, *T_NR_*, was longer than the allowed RT, participants would fail to cancel the impending response. This led to a predicted probability of being correct that increased smoothly as a function of allowed RT, as observed in the data. An analogous model was applied to the NR-to-R condition. In this case, the commitment to initiating a response was assumed to be made at a random time 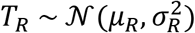 after the circle changed color and, on a given trial, they would succeed at correctly generating the response at the right time only if *T_R_* is shorter than the allowed RT on that trial. Thus, *μ_NR_* and *μ_R_* represented the average times required either to cancel a movement or initiate a movement, respectively.

We fitted these models to each participants’ data via maximum likelihood estimation, yielding model fits that closely matched the empirical data (Fig. 2D, Solid lines; Fig. S1). By comparing the estimates of *μ_NR_* to *μ_R_* across participants, we found that the time required to cancel the initiation of an impending response (*μ_NR_*: 294.5 ms ± 28.3 ms; mean ± s.d.) and the time required to initiate a response (*μ_R_*: 289.0 ms ± 24.4 ms; mean ± s.d.) were not significantly different from one another (*μ_nr_ – μ_r_*: 5.5 ms ± 24.4 ms, Cohen’s d = 0.207, t34 = 1.34, paired t-test, p = 0.19, 95% CI: [-2.8ms, 13.9ms]; Fig. 2E) and were also highly correlated across participants (ρ = 0.58, p < 0.001). The equivalence and noninferiority test demonstrated that these two speeds were statistically equivalent (p_lower_ < 0.001; p_upper_ = 0.004, 90% CI: [-1.5ms, 12.5ms]). Thus, when we compared action initiation and action inhibition under experimental conditions that were as closely matched as possible, we found that generating a response takes the same amount of time as canceling one.

Because *μ_NR_* and *μ_R_* represent the RT at which performance accuracy reaches the center of the speed–accuracy trade-off function, their estimates were sensitive to exactly how we defined whether a trial was performed correctly or not. In our initial analysis, a late response (if the circle was below the target line without overlapping it at the time the response was made) of an NR-to-R trial was considered as incorrect, whereas in R-to-NR trials, it was impossible to distinguish between trials in which a response was cancelled at the correct time, and trials in which a response was delayed before being cancelled. This unavoidably imposed a stronger requirement of timing precision on initiating a response than cancelling a response (asymptotic accuracy: β = 0.89 ± 0.06 vs. β = 0.96 ± 0.05; t34 = −5.26, paired t-test, p < 0.0001, 95% CI: [−0.09, −0.04]). However, this aspect of the analysis did not greatly bias our estimate of the time course of action initiation. When we relaxed the timing requirement for correct responding in the NR-to-R condition to match the asymptotic accuracy in the R-to-NR condition (0.94 ± 0.06 vs. 0.96 ± 0.05; t34 = −1. 9, paired t-test, p = 0.07, 95% CI: [-0.04ms, 0.01ms]), the resulting estimate of *μ_R_* (286.3 ms ± 23.6 ms) was only 2.7 ms different from the original analysis and was only 8.2 ms shorter than *μ_NR_* (*μ_NR_ – μ_R_*: 8.2 ms ± 23.5 ms, Cohen’s d = 0.31, t34 = 2.06, paired t-test, p = 0.047, 95% CI: [0.0ms, 16.3ms], power = 0.5; Fig. S3). Furthermore, the difference between *μ_NR_* and the updated *μ_R_* fell within the equivalence bound between −16.5 ms and 16.5 ms (p_lower_ < 0.001; p_upper_ = 0.02, 90% CI: [1.5ms, 14.9ms]). Therefore, our conclusion about action initiation and action inhibition having a similar time course was not strongly biased by the presence of late responses.

Similar to other analyses of behavior in stop-signal tasks, the preceding analyses relied on designating a proxy RT for trials where a response was never generated and it is possible that this might have influenced the results. To avoid this issue, we considered an alternative analysis of behavior in the NR-to-R condition based only on trials in which a response was generated and the actual time of those responses as a function of the intended allowed RT (i.e., time interval between color change and the target line). We observed that, when the intended allowed RT was very short (e.g., < 200 ms) participants often did generate a response, but much later than required by the task. This suggested that they switched to performing the task in a reactive mode, responding as soon as they could after the color change, even if this meant their response would be late (Fig. 3A). We assumed that the timing of this switch of strategy coincided with the minimum time at which participants could successfully initiate a movement in response to the color change. To estimate the time of this switch, *μ_T_*, we fit a simple model of the timing of participants’ responses as a function of intended allowed RT (Methods). The model comprised two linear components: one to represent accurately timed responses at longer allowed RTs, and one to represent delayed, reactively generated responses at very short allowed RTs (Fig. 3A; Solid light blue line), with the transition between these occurring at *μ_T_*. The estimates of *μ_T_* (302.3 ms ± 39.0 ms; mean ± s.d.) were in close agreement with the original estimate of *μ_R_* based on designative proxy RTs (ρ = 0.65; p < 0.0001). These alternative estimates of the time needed to initiate a movement also did not differ significantly from the time that participants needed to cancel a previously intended response, *μ_NR_* estimated in the R-to-NR condition (*μ_NR_ – μ_T_*: −7.7 ms ± 37.7 ms, Cohen’s d = −0.22, t34 = −1.25, paired t-test, p = 0.218, 95% CI: [-20.3ms, 4.8ms]; Fig. 3B). Results from the equivalence and noninferiority tests confirmed that these two speeds were similar to each other (p_lower_ = 0.003; p_upper_ < 0.001, 90% CI: [−18.4ms, 2.8ms]). This result accords with our initial analysis that action initiation and action inhibition follow a similar time course.

**Fig. 3.**
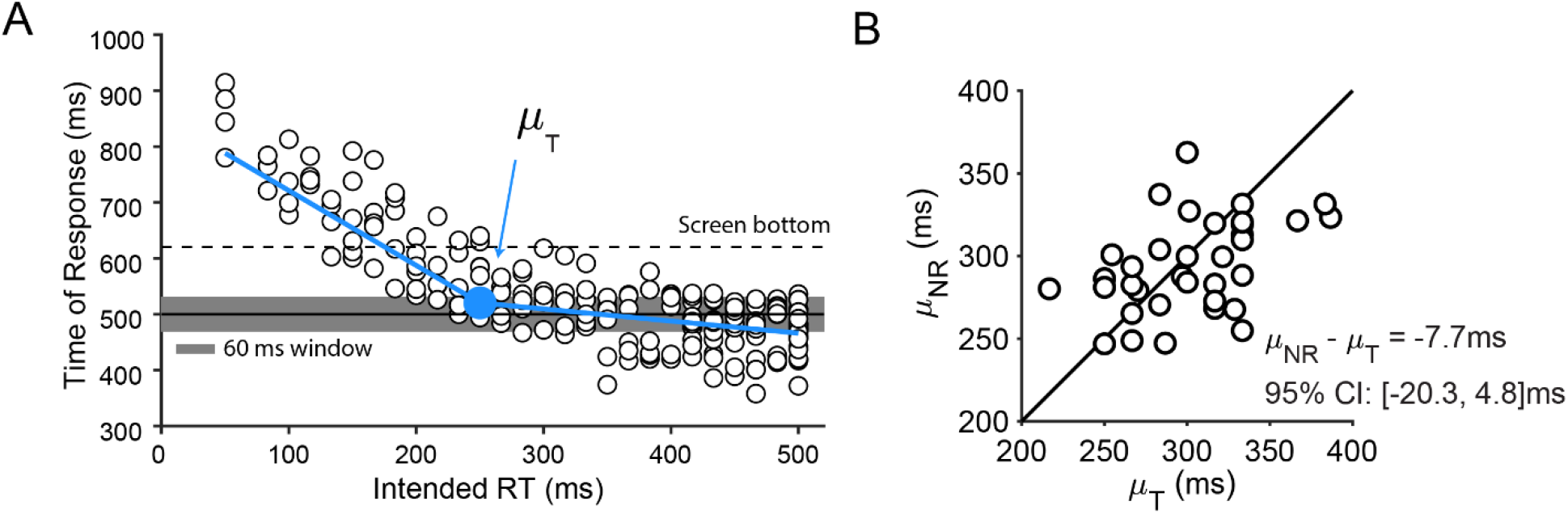
Alternative analysis based on the pattern of delayed responses. A) The time of response data as a function of intended RT from the same exemplar participant as in Fig. 2B. The time of response was measured as the time interval between trial start and the time at which a response is made. Intended allowed RT was the time interval between color change and the target line. Since this analysis focused only on trials in which a response was actually generated, it did not rely on approximating unobserved RT. We fitted the time of response data with two linear functions of intended allowed RT, which intersected at *μ_T_* – a parameter that represents the minimum allowed RT at which an accurately timed response could still be made. B) Across all participants, the estimates of *μ_T_* (302.3 ms ± 39.0 ms; mean ± s.d.) – the alternative estimate of time required to initiate a response – were not significantly different from *μ_NR_* (294.5 ms ± 28.3 ms; mean ± s.d.), the original estimate of the time required to cancel a response.

## Discussion

By matching experimental conditions as closely as possible between initiating an action and inhibiting an impending action, we found that the time requirements for these two processes are comparable, both at around 290 ms (note that this included a ~33 ms delay caused by the visual display, which was not accounted for in our calculations). Our findings demonstrate that the speed disadvantage often observed in action initiation is not because it is inherently slower than action inhibition. Rather, the fact that action inhibition is usually faster than action initiation is attributable to asymmetries in urgency for these processes in the conventional stop-signal paradigm.

Unlike the dedicated effort to understanding action inhibition over the past decades (Hannah and Aron, 2021; Matzke et al., 2018; Verbruggen and Logan, 2017), action initiation has received much less attention and is less comprehensively understood. One possible reason why there has been such a strong focus on action inhibition is because it has been thought to be a crucial component of executive control (Aron et al., 2014; Logan and Cowan, 1984; Verbruggen and Logan, 2008), while action initiation is considered to be a more elementary process and of less-critical importance for cognitive control. In real life circumstances, however, making a response that is not planned to be generated at the first place (i.e., action initiation) can be just as important as stopping a response that is about to be initiated (i.e., action inhibition). For instance, a common example of action inhibition is that a pedestrian about to step into the street may need to swiftly abort this act to avoid a fast-approaching car. However, if they realize that their bus is about to leave, they may need to hastily cross the street. We therefore argue that the decision to act, from a state of inaction, may be just as crucial component of executive control as the converse process of action inhibition.

The speed of generating a movement has often been measured in simple reaction time tasks, a research field that take place in isolation from the research of action inhibition. However, typical simple reaction times identified in these studies (Luce, 1991; Welford, 1980) are roughly comparable to the time to inhibit an action (He et al., 2021; Leunissen et al., 2017; Logan and Cowan, 1984; Matzke et al., 2021), with both typically around 200 – 250 ms. Furthermore, both times can be reduced to around 150 ms if triggered by an unexpected event (Carlsen et al., 2004; Haith et al., 2016; Wessel and Aron, 2017). This evidence is consistent with our findings. One major difference between the simple reaction time task and our experiment is that, in simple reaction time tasks, all trials demand a response to be made, while in our NR-to-R condition where the speed of action initiation was measured, most trials did not require a response. It is not yet known whether the percentage of trials that require a response (or no response) would systematically affect the speed of action initiation and whether such an effect is accordance with its role on action inhibition (Aron, 2011; Dunovan et al., 2015).

Because inhibition works on an action that is about to be initiated, our ability to stop an action depends strongly on the time course of action initiation. The fact that action initiation and action inhibition are equally fast suggests that in order for stopping a response to be successful on a regular basis, action initiation would have to be systematically delayed. Indeed, many conventional stop-signal tasks are deliberately designed to slow movement initiation by, for example, including a choice between possible actions, in order to promote successful inhibition. However, slowing of responses has been widely observed in previous stop-signal tasks over and above that imposed by task design (e.g., Corneil et al., 2013; Gulberti et al., 2014; Lappin and Eriksen, 1966; Leotti and Wager, 2010; Özyurt et al., 2003; Verbruggen and Logan, 2008). As a result, response times in stop-signal tasks are typically prolonged (e.g., Leunissen et al., 2017) and are accompanied by delayed motor cortex excitability (Rawji et al., 2022). Intriguingly, it has been found that participants often delayed the response by an amount slightly greater (~ 5 to 20 ms) than the time lag between the primary go and stop signals (Lappin and Eriksen, 1966), which is just enough to allow them to stop a response (it also suggests that inhibition is as fast as initiation). This flexibility in deciding when to initiate a movement is related to the concept of “freedom for immediacy” – the capacity to decouple our responses to external stimuli from the appearance of the stimulus itself (Haggard, 2008; Haith et al., 2016; also, Kornblum, 1973; Ollman and Billington, 1972). The critical role of flexibility in movement initiation is exemplified in the stop-signal task, as being able to successfully stop an action depends not only on the response inhibition process being fast (Boucher et al., 2007; Dunovan et al., 2015; Logan et al., 2015; but see, Salinas and Stanford, 2013), but can also result from slowed initiation in response to the go cue. The flexibility of action initiation is therefore just as integral to behavior in stop-signal tasks as action inhibition and might be considered an equally important component of executive control.

The race model is widely used to study the relationship between action initiation and action inhibition (Logan and Cowan, 1984). However, the “go” process that determines a response to be made often consists of multiple distinct aspects of action control (Hanes and Carpenter, 1999; Logan et al., 2014). The fact that delaying the time to initiate an action is an essential part of stopping behavior implies that the “go” process includes the decision about whether to respond and when to respond. In tasks imposing multiple actions (e.g., Verbruggen et al., 2019), a process of deciding what response to prepare is also needed. For example, the go process in the classic stop-signal task would be different from the go process in our task in which a response is required to be made at a pre-determined time (see also, Dunovan et al., 2015). Previous research has tended to assume that action initiation directly follows and is triggered by action preparation (Donders, 1969; Sternberg, 1969). However, recent neural and behavioral evidence in motor control has stressed that these processes are very likely mechanistically independent (Ames et al., 2019; Elsayed et al., 2016; Haith et al., 2016; Haith and Bestmann, 2020; Kaufman et al., 2016; Lara et al., 2018). For example, it may not be necessary for a go process to reach a fixed threshold level before a response is initiated (Jagadisan and Gandhi, 2017). We suggest that future studies may incorporate action preparation and action initiation as distinct processes. This may help us determine how action initiation and action inhibition work together. For example, when we inhibit a response, what aspect of the response is stopped (e.g., a “prepared” response or an “already-initiated” response) and how they are stopped (e.g., Boucher et al., 2007; Dunovan et al., 2015; Logan et al., 2015; Salinas and Stanford, 2013).

By constraining the decision about when to act as much as possible, as well as measuring action initiation and action inhibition under two separate and symmetric conditions, we observed comparable and highly correlated speeds between these two processes. This finding raises the possibility that action initiation and action inhibition might not reflect two distinct processes, as is typically assumed, but may in fact reflect a single underlying process. Under this view, action initiation and action inhibition may be two opposing states of a single process reflecting the decision about whether to act or not. Indeed, in the context of voluntary movement, the decision of whether or not to act is thought to be an important function that is independent of decisions about how and when to act (Brass and Haggard, 2008).

Action inhibition in humans is thought to be controlled through a particular prefrontal-cortex – basal-ganglia hyperdirect pathway. This pathway is believed to serve as an “emergency brake” that can abruptly abort a response, whose initiation is governed through a direct pathway (Aron, 2007; Aron et al., 2014; Chen et al., 2020; Dunovan and Verstynen, 2016; Hannah and Aron, 2021; Jahanshahi et al., 2015; Wessel and Aron, 2017; Wiecki and Frank, 2013). However, this understanding has largely been established through the lens of conventional stop-signal tasks. If, however, it really is the case that action initiation and action inhibition are more closely related than previously thought, the fundamental role of the neural mechanisms thought to be dedicated to action inhibition may need to be revisited. These mechanisms might also be important for action initiation, which however cannot be established without directly contrasting action inhibition and action initiation. Indeed, it has been reported that in some cases, circuits thought to be important for action inhibition have been found to also be engaged during action selection and production (Filevich et al., 2012; Mostofsky and Simmonds, 2008). We suggest that, to better understand the neural basis of action initiation and action inhibition, it is critical to examine them in tandem, under closely matched experimental conditions as we have shown here.

## Methods

### Participants

Thirty-six right-handed participants (15 female; 1 non-binary) between 18 and 41 years of age took part in the study. The experimental procedure was approved by the Johns Hopkins School of Medicine Institutional Review Board. All participants gave written informed consent and received $15 per hour for their participation. Data from one participant who did not follow the task instructions well were excluded from analyses (Fig. S1 and S2).

### General procedures

Participants sat in front of a laptop with a gray screen and with a key pad next to it. The key pad was positioned so that participants could comfortably rest the index finger of their right hand on a mechanical key mounted on the key pad. On every trial, a white target line was placed with the same distance from the bottom of the screen and a circle was displayed at the top center of the screen (Fig. 1). Once the trial started, the circle moved downwards vertically and participants were asked to either press the key or do nothing when the moving circle reached the target line. The circle stopped moving once a response was registered, or it kept moving toward the bottom of the screen.

For trials where a response was required, a red cross mark was shown if the circle did not intersect the line when it was stopped by a response or if the circle left the screen with no response having been generated, while a green check mark appeared if the circle intersected the target line when it was stopped by a response. The feedback was used to encourage participants to respond with accurate timing and minimize tendencies to delay their response in order to gain more time to make decisions. In trials in which no response was required, a green check mark was displayed if the circle left the screen without a response having been made, while a red cross was displayed if any response was generated at all. Whether or not a response was required in a given trial depended on the color of the circle (white or black). Since perceptual processing plays a critical role in motor response inhibition (Salinas and Stanford, 2013), we counterbalanced the association between response and the color of the moving stimulus so that for half of the participants, white color cued a response (i.e., press the key) and black color indicated that no response was needed (i.e., do not press the key), while this association was reversed for the other half participants, so as to control for the potential perceptual differences between black and white colors.

### Criterion task

Before the experimental trials began, participants completed two criterion blocks. In these two blocks, all trials required a response in order for participants to become familiar with the timing requirement of the response. The meaning of the color used in this task was consistent with that used in subsequent tasks for each individual. In the first and easier criterion block, the moving circle started from the top center of the screen and dropped toward the bottom of the screen with a constant speed, which took 900 ms in total. A white target line was placed 750 ms from the top and thus 150 ms from the bottom. The circle diameter was sized such that it took 120 ms for the circle to move across the target line. The block ended with five consecutive correct responses (i.e., any part of the circle stopped on the line). Participants then performed the second and more difficult criterion block which matched the conditions of the main experiment, i.e., the diameter of the circle was reduced to 60 ms and it took 500 ms from the trial onset to the center of the circle intersecting the target line and another 120 ms to the bottom of the screen. Similarly, 5 consecutive correct trials were required to end this block.

After successfully completing these two criterion blocks, participants then performed the main task with a response-to-no response (R-to-NR) condition and a no response-to-response (NR-to-R) condition, the order of which were counterbalanced across participants. Each condition consisted of 6 blocks and each block had 100 trials.

### R-to-NR Condition

This task is also known as the adaptive stop-signal task (Coxon et al., 2006; Leunissen et al., 2017; Slater-Hammel, 1960) and has been used to examine how fast participants can decide to cancel a prepared response that was originally planned to be executed. The moving circle always started with the color that cued a response (white for half of participants and black for the other half). In a random ~30% of trials (204 out of 600 trials), the circle turned to the not-responding color while it was moving towards the target line. The time of color switch before the center of the circle intersected the target line was randomly drawn from a uniform distribution between 50ms to 500 ms with a step size of 16.7ms. The choice of this step size was constrained by the refresh rate of the monitor, which was 60 Hz. Thus, there were 28 possible time points at which the circle color changed. The closer the time point was from the targe line, the shorter time available to make a decision.

### NR-to-R Condition

This task, conceptually similar to the timed-response task commonly used in motor reaching task (Ghez et al., 1997; Haith et al., 2016), was used to examine how fast participant can initiate a response. Trials started with the circle defaulted to the not-responding color (white for half of participants and black for the other half) and switched to the responding color in a random subset of trials. Consistent with the R-to-NR condition, the proportion of color-switch trials was ~30% of trials (204 out of 600 trials) and the time of color change ranged from 50 to 500 ms. These switch trials and their corresponding color change times were matched between these two conditions on a trial-by-trial basis. One participant showed clear evidence of guessing the required response in the NR-to-R condition (Figs. S1 and S2) and we therefore excluded this participant from subsequent analysis.

## Data analysis

### speed–accuracy trade-off

Our primary analysis was focused on the trials in which the color switched in both conditions. We assessed the time course over which participants were able to abort an impending action in the R-to-NR condition by constructing a speed–accuracy trade-off relating the time available to cancel a response and the probability of the response being successfully cancelled. For visualization purposes, we estimated this speed–accuracy trade-off using a 50 ms sliding window on the time available to cancel a response (i.e., allowed RT, see below for details). Similarly, the speed–accuracy trade-off can also reveal how rapidly a response can be initiated in the NR-to-R condition.

### Response correctness

In our original speed–accuracy trade-off analysis (Fig. 2), a trial in which a response was required was considered to be successful only if a response was made while the circle was not below the target line (i.e., within a −30 ms time window below the target line). In practice, participants often did generate a response, but did so after the circle was no longer overlapping the line. We designated these trials as failures and considered them to be equivalent to not generating a response at all. However, we designated a trial that does not require a response as correct if participants did not press the button before the trial ended. This non-produced response may be more than 30 ms after the target line if it were generated.

This designation of correctness included a strong requirement of timing accuracy when initiating a response in the NR-to-R condition but not when cancelling a response in the R-to-NR condition (Fig. 2C). This asymmetry in the analysis may have affected our estimation of the relative timing of going and stopping. To match the timing requirement, we further used a −35 ms time window below the target line. Results from these two different criterions for correctness are consistent with one another (Fig. S3 and Main Text).

### Allowed RT

By manipulating the time at which circle changed its color (between 50 ms and 500 ms before the targeted line), we forced participants to cancel an intended response in the R-to-NR condition or press a button in the NR-to-R condition within a particular amount of time, referred to as allowed RT. In the R-to-NR condition, when participants failed to cancel an impending response, the allowed RT was quantified as the time elapsed from color change to the time of the button press. When no response was generated, the actual allowed RT was not observable and so instead the allowed RT was approximated by the intended allowed RT, i.e., the time interval between the center of the circle and the target line at the moment of color change. In the NR-to-R condition, the allowed RT was calculated as the time interval between color change and the time of response if participants pressed a button before or when the circle reached the target line, whereas it was approximated as the interval between color change and the target line (i.e., intended allowed RT) if participants did not generate a response or the response was made later than the target line (i.e., the circle was no longer overlapping the target line).

When using the intended RT described above, we inherently assumed that the not-produced response would have had accurate timing, if it was produced, which, however, is not true in reality. All participants had an idiosyncratic tendency to respond consistently earlier or later than the target line (Fig. S4). To better approximate the true RT, we first calculated how much later or earlier each participant responded to trials in non-switch trials in which the circle was in the responding color throughout the trial (i.e., 396 out of 600 trials, about 70% in the R-to-NR condition). From these measurements, we randomly drew a sample and added it to an intended RT to approximate the unobservable true allowed RT. We repeated this bootstrapping process 1000 times for each individual and modelled the mean speed–accuracy trade-off.

### Modelling speed–accuracy trade-off

To quantify the speed–accuracy trade-off, we assumed that cancelling an intended response in the R-to-NR condition occurred at a random time 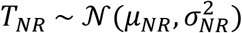. A response would be correctly aborted with a probability *α_NR_* (close to 0) if the available allowed RT was shorter than *T_NR_*. Similarly, a response would be correctly aborted with probability *β_NR_* (close to 1) if allowed RT was long than *T_NR_·β_NR_* that is less than 1 indicates that a response may not be correctly stopped even given long allowed RT, reflecting ‘trigger failure’ where the stop process is never initiated due to participants being inattentive to the signal. So unlike classic stop signal studies in which the estimate of the stopping speed may be biased by trigger failure, (Band et al., 2003; Matzke et al., 2017; Verbruggen et al., 2019), our model takes such effect into account to estimate the stopping speed *μ_NR_*.

According to our model, the probability, in trial *i*, of observing a correct response cancellation (c = 1), given the preparation time (*t^i^*) is given by:

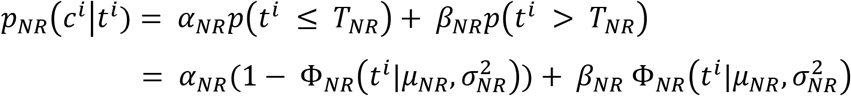

where 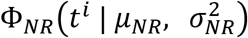 is the cumulative normal distribution of *T_NR_*.

Similarly, in the NR-to-R condition, the probability of correctly initiating a response given the preparation time (*t^i^*) is:

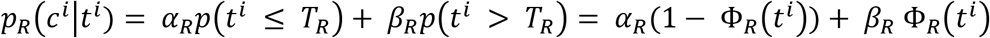

where 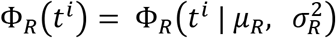 is the cumulative normal distribution of *T_R_*.

We estimated the parameters using maximum likelihood estimation with the MATLAB function fmincon.

### Modelling time of response

Time of response was the time elapsed from the trial onset to the time at which the response was made, if any. In our speed–accuracy trade-off analysis, we relied on a proxy RT for trials in which a response was not produced. To avoid this reliance, we also estimated the speed of making a response in the NR-to-R condition by fitting the time of response, y, with two linear functions, which intersected at an intended allowed RT of *μ_T_*:

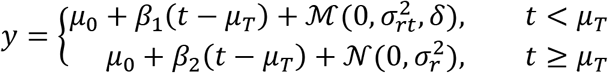

In this model, we took the time of transition between these components, *μ_T_*, as the minimum time at which participants could initiate a response on time following the color change. At *μ_T_*, a response needed to be made without any delay so that it could land on the target line. *μ_0_* is the time of response when *t* = *μ_T_*. We assumed that for *t* < *μ_T_*, participants behaved in a reactive manner to the appearance of stimulus and that, therefore, the time of response would follow a typical reaction time distribution. In particular, we observed that some participants occasionally generated times of response that were longer than most of their responses, so we assumed that the residual term in the upper equation followed an exponentially modified Gaussian distribution 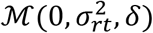. This choice did not lose its generality if participants did not generate some uncommon late responses, because when *δ* is close to zero, 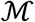 approaches to be a Gaussian distribution. When *t* ≥ *μ_T_*, participants would time the response and press the button at around the target line. Thus, we assumed that residual term of the lower equation in this case followed a Gaussian distribution 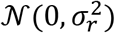. Parameters *μ*_0_, *β*_1_, *β*_2_, *μ_T_*, *σ_rt_*, *σ_r_*, *δ*, were estimated by maximum likelihood estimation with MATLAB function fmincon. To avoid a local minimum estimation, we ran the maximum likelihood estimation with 100 random starting values. A parameter recovery analysis indicated that our model fitting yielded unreliable estimation of true parameters (Fig. S5). We found, based on parameter recovery, that it was better to constrain *σ_rt_* to be greater than 0 (lower bound of 0.005), in order to avoid poot quality fits. For the same reason, we also regularized the fits by penalizing the log-likelihood with:

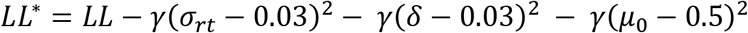

*σ_rt_* and *δ* were included to avoid unrealistic estimation of *σ_rt_* ≈ 0 and *δ* ≈ 0 (Fig. S5). We chose 0.03 for *σ_rt_* and *δ* as it was the mean value of initial estimation across participants. We also regularized *μ*_0_ and set it to 0.5 s because our data showed that participants tended to respond around the target line given a long enough RT (Fig. S4). We set *γ* = 2000, which avoided overfitting these three parameters to the particular value we selected. Parameter recovery demonstrated that this regularized fitting procedure led to reliable estimation of the true parameters when applied to synthetic data (Fig. S6).

### Statistical analysis

Data (e.g., *μ_NR_* vs. *μ_R_*) were analyzed using paired t-test at the significant level of *α* = 0.05 after examining the normality of samples. Because non-significant outcomes from hypothesis testing do not necessarily support that two samples are not different from one another, we further conducted the equivalence and noninferiority test (Lakens, 2017; Walker and Nowacki, 2011). Power analysis indicates that the sample size n = 35 (out of 36) had 80% power to detect an effect size of 0.7 between conditions (Lakens, 2017). Therefore, we set the upper and lower bounds of the equivalence and noninferiority test as 0.7 and −0.7, corresponding to the equivalence bound between 16.5 ms and −16.5 ms in the scale of reaction time.

## Author Contributions

Y.D., A.D.F., and A.M.H. conceptualized the experiment; A.D.F. programmed the task; Y.D., A.D.F., and D.M.M. collected data; Y.D. performed data and statistical analyses; Y.D. prepared the figures; Y.D. drafted the manuscript; Y.D., A.D.F., D.M.M., and A.M.H. revised the manuscript and approved final version of the manuscript.

## Data Availability

All data generated from this study and the code for reproducing the experiment are available at the corresponding author’s personal github page: https://github.com/YueDu-Science/Inhibition-Initiation. The data will also be available at https://archive.data.jhu.edu/. The code for reproducing the results will be available at https://github.com/YueDu-Science/Inhibition-Initiation. We are currently working on cleaning and organizing the code.

## Supplement Figures

**Fig. S1.**
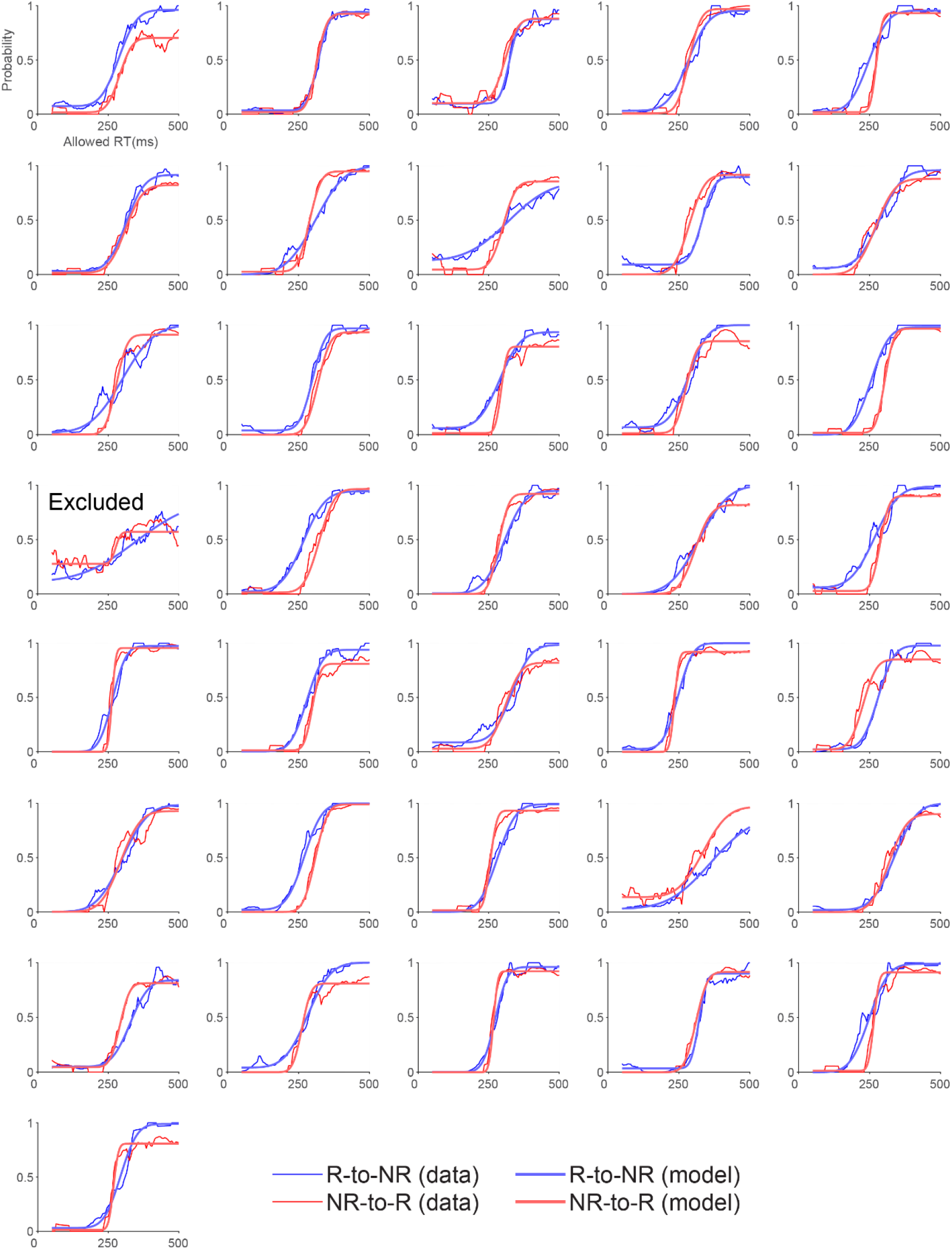
Speed–accuracy trade-off for each individual. Most participants’ performance (thinner lines) was similar to the exemplar participant shown in Fig. 2. The accuracy was close to zero when allowed RT was very short (i.e., < 100 ms) and it increased with longer allowed RT. However, one participant exhibited abnormally high accuracy even when allowed RT was less than 100 ms, suggesting that this particular participant randomly guessed whether to initiate or cancel a response instead of following the instruction for each condition. We excluded this participant from further analysis.

**Fig. S2.**
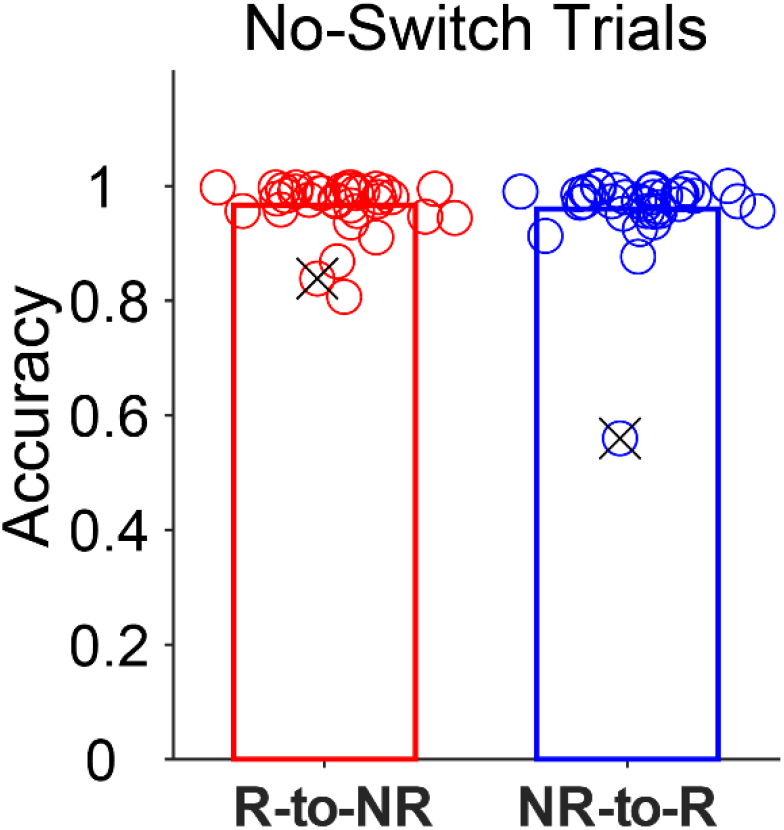
Response correctness in trials that stimulus circle did not change color. The high accuracy in no-switch trials (i.e., Making a response in the R-to-NR condition or not making a response in the NR-to-R condition) revealed that participants did not behave randomly except one participant, who exhibited a chance-level performance in the NR-to-R condition. This participant was the one who also produced higher accuracy in switch trials even when allowed RT was very short shown in Fig. S1. Since we aimed to compared the individual-wise performance between two tasks, this participant’s data from both conditions (crossed circles) were excluded for our analyses reported in main texts.

**Fig. S3.**
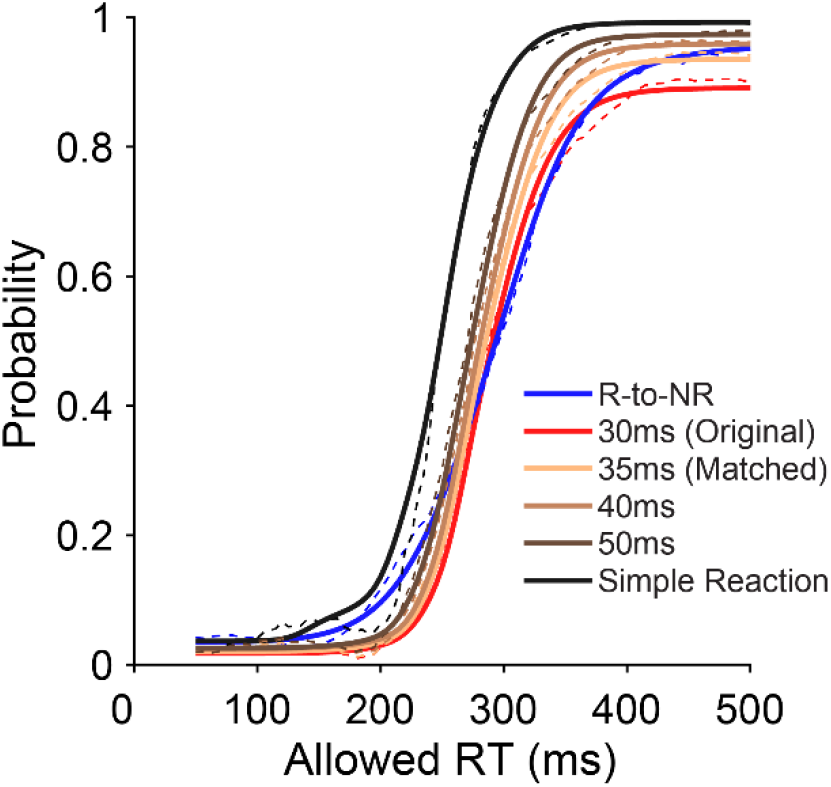
The effect of correctness criterion on the estimated speed of response initiation. A) The shape of the speed–accuracy trade-off in the NR-to-R condition depended on how we defined a correct responding trial. The original tolerance was 30ms (i.e., a half size of the stimulus circle). Correctness defined on this small-time window did not include trials where a response was made 30 ms or more later below the target line, resulting in a lower accuracy in the NR-to-R condition than the R-to-NR condition when RT was long enough (red vs. blue). To match accuracy rates across conditions, we instead used a timing tolerance of 35 ms that yielded a similar accuracy level between conditions for trials with long RT (light brown). In this case, we still found that the timing of response cancellation was not different from the timing of response generation (blue vs. light brown). In addition, the original estimation of response initiation speed was around 285 ms. This slow speed was, at least partially, caused by the timing requirement of initiating a response on time in the NR-to-R condition. Further broadening timing tolerances from 35 ms to 50 ms led to superior speed–accuracy trade-offs with faster mean speeds of responding than the original (brown lines). The speed was even faster (248.7 ms ± 31.7 ms) when all trials with a response at any time were considered correct (the ‘simple reaction’ mode; black line). Solid lines: model fitting; Dashed lines: data.

**Fig. S4.**
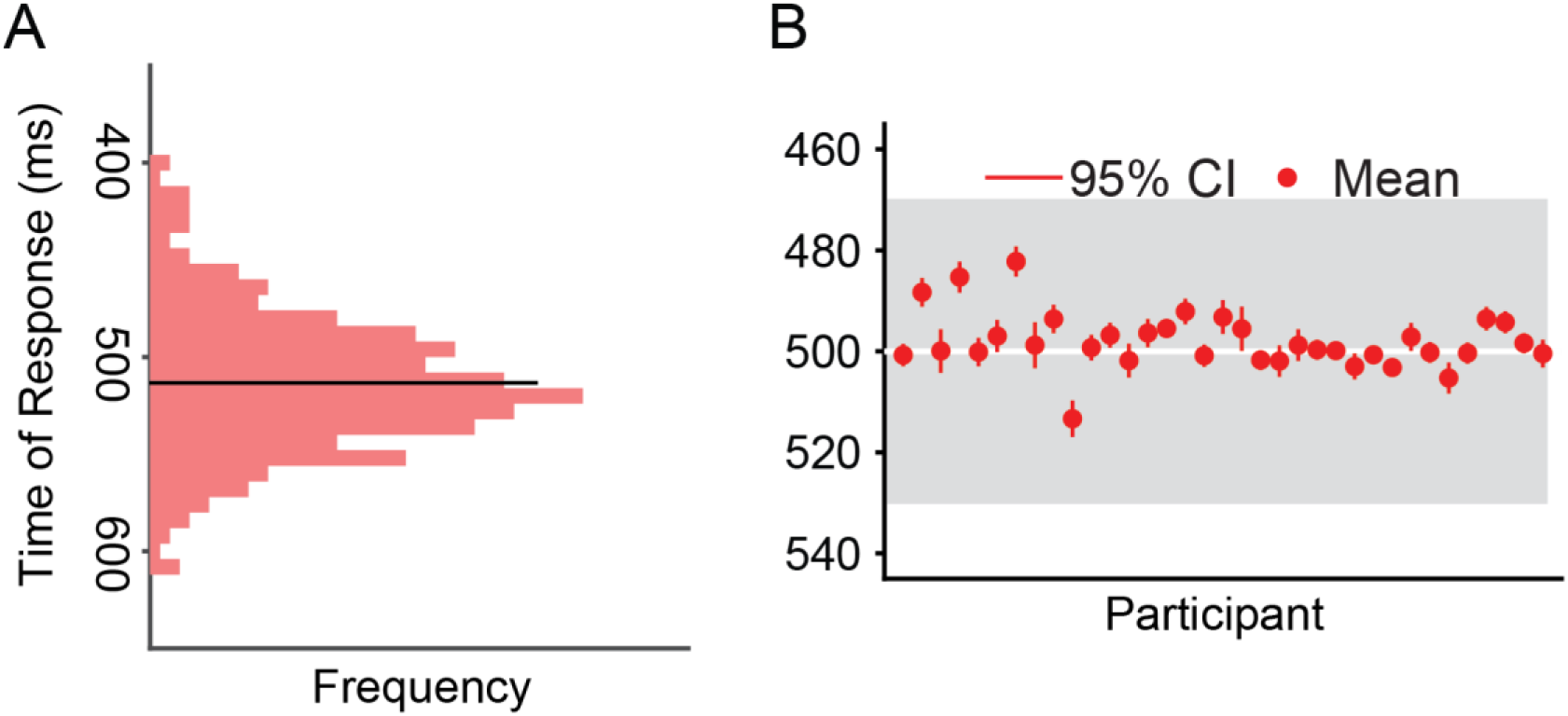
Idiosyncratic tendencies to respond consistently earlier or later than the target line. A) Data from the same exemplar participant as in Fig. 2B. The time of response, measured for trials that the circle started and stayed as the same color that required a response in the R-to-NR condition, was calculated as the time interval between trial onset and the time at which a response was made. This participant tended to respond consistently slightly later than the target line (i.e., 500 ms). B) The mean time of response for each individual participant. All participants had an idiosyncratic tendency to respond consistently earlier or later than the target line, although the circle still overlapped the target line when they responded (grey area representing the diameter of 60 ms of the circle).

**Fig. S5.**
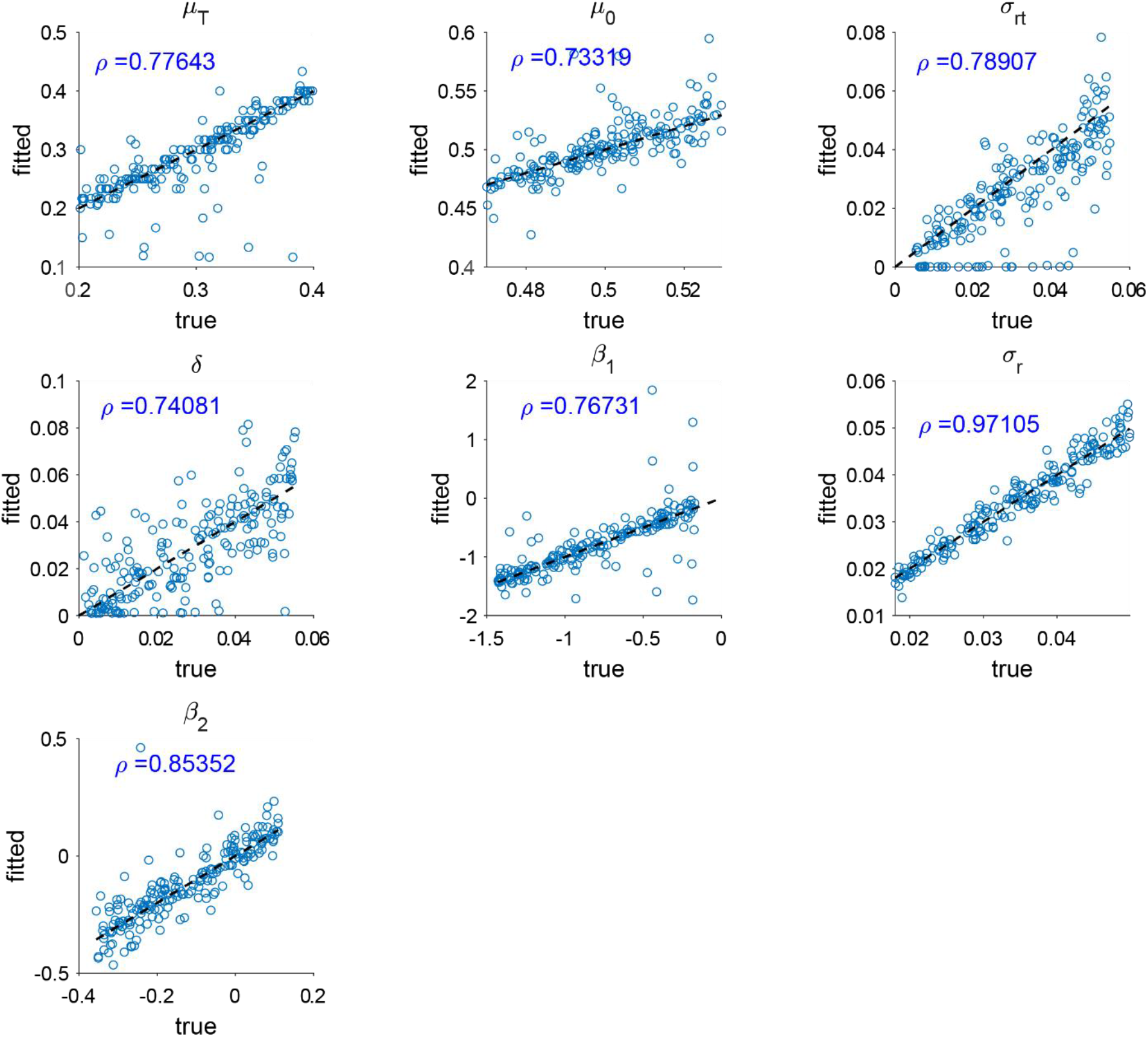
Parameter recovery of the model used to fit time of response data without regularization terms. We used the model to generated synthetic datasets matching the amount of data collected from each participant. We generated datasets based on a range of true underlying parameter values and then used maximum likelihood estimation to try to recover the true underlying parameters. Each panel shows a different parameter (unit of measurement: second), with true value used in the simulation on the x-axis and the estimated value on the y-axis. For most parameters, including the key parameter of interest, *μ_T_*, parameter recovery was not accurate.

**Fig. S6.**
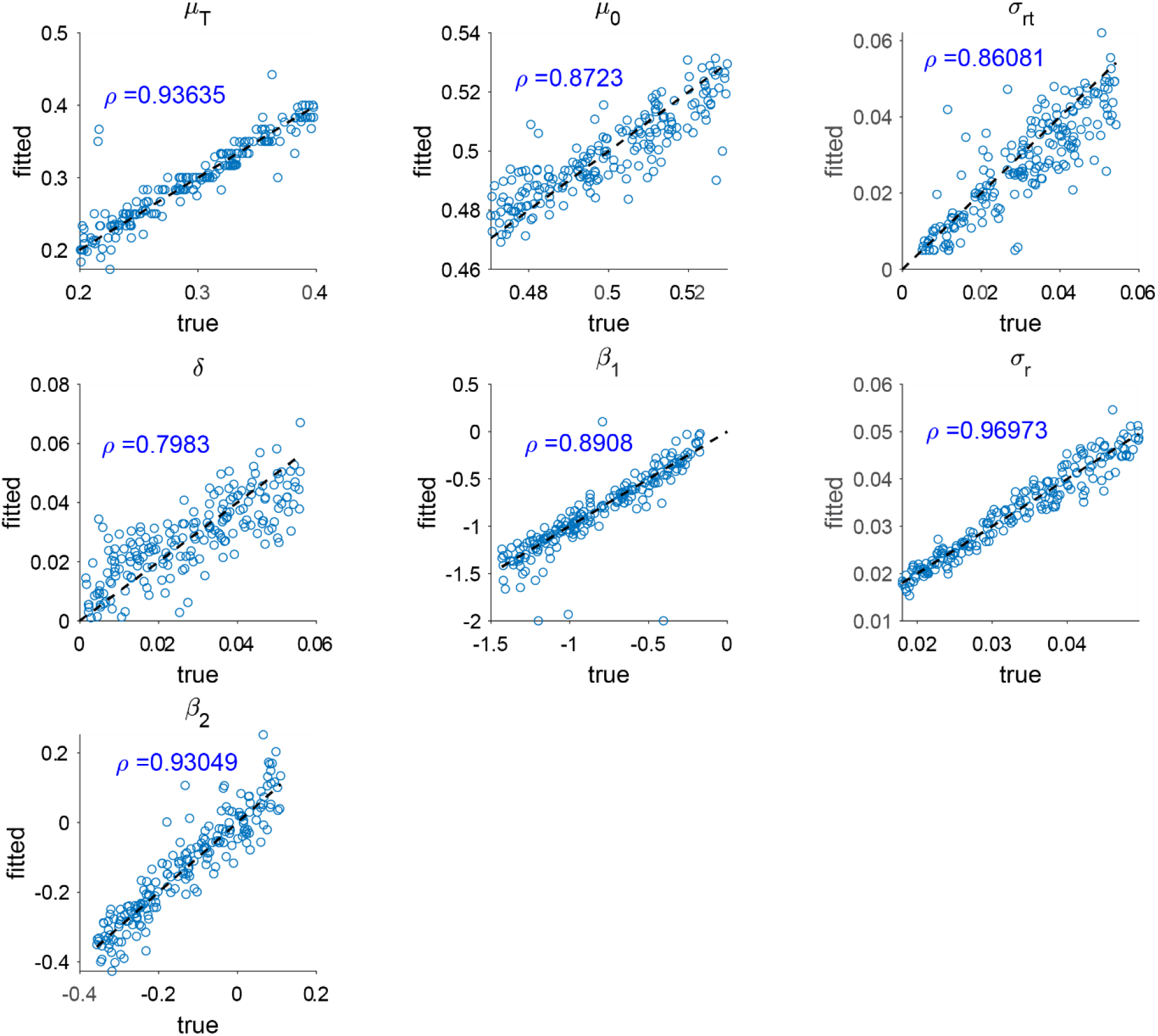
Parameter recovery of the model used to fit time of response data with regularization terms. After adding regularization terms with respect to *μ*_0_, *σ_rt_*, and *δ*, the reliability of the parameter estimation became notably improved. The correlation between true values and estimated values of *μ_T_* = 0.94, which is much higher than the corresponding value of 0.78 in the model without regularization (Fig. S5).

## References

Ames KC, Ryu SI, Shenoy KV. 2019. Simultaneous motor preparation and execution in a last-moment reach correction task. Nature communications 10:1–13.

Aron AR. 2011. From reactive to proactive and selective control: developing a richer model for stopping inappropriate responses. Biological psychiatry 69:e55–e68.

Aron AR. 2007. The neural basis of inhibition in cognitive control. The neuroscientist 13:214–228.

Aron AR, Robbins TW, Poldrack RA. 2014. Inhibition and the right inferior frontal cortex: one decade on. Trends in cognitive sciences 18:177–185.

Band GP, Van Der Molen MW, Logan GD. 2003. Horse-race model simulations of the stop-signal procedure. Acta psychologica 112:105–142.

Bissett PG, Logan GD. 2014. Selective stopping? Maybe not. Journal of Experimental Psychology: General 143:455.

Boucher L, Palmeri TJ, Logan GD, Schall JD. 2007. Inhibitory control in mind and brain: an interactive race model of countermanding saccades. Psychological review 114:376–397.

Brass M, Haggard P. 2008. The what, when, whether model of intentional action. The Neuroscientist 14:319–325.

Carlsen AN, Chua R, Inglis JT, Sanderson DJ, Franks IM. 2004. Prepared movements are elicited early by startle. Journal of motor behavior 36:253–264.

Chen W, de Hemptinne C, Miller AM, Leibbrand M, Little SJ, Lim DA, Larson PS, Starr PA. 2020. Prefrontal-subthalamic hyperdirect pathway modulates movement inhibition in humans. Neuron 106:579–588.

Corneil BD, Cheng JC, Goonetilleke SC. 2013. Dynamic and opposing adjustment of movement cancellation and generation in an oculomotor countermanding task. Journal of Neuroscience 33:9975–9984.

Coxon JP, Stinear CM, Byblow WD. 2006. Intracortical inhibition during volitional inhibition of prepared action. Journal of neurophysiology 95:3371–3383.

Donders FC. 1969. On the speed of mental processes. Acta psychologica 30:412–431.

Dunovan K, Lynch B, Molesworth T, Verstynen T. 2015. Competing basal ganglia pathways determine the difference between stopping and deciding not to go. Elife 4:e08723.

Dunovan K, Verstynen T. 2016. Believer-skeptic meets actor-critic: rethinking the role of basal ganglia pathways during decision-making and reinforcement learning. Frontiers in neuroscience 10:106.

Elsayed GF, Lara AH, Kaufman MT, Churchland MM, Cunningham JP. 2016. Reorganization between preparatory and movement population responses in motor cortex. Nature communications 7:1–15.

Filevich E, Kühn S, Haggard P. 2012. Negative motor phenomena in cortical stimulation: implications for inhibitory control of human action. cortex 48:1251–1261.

Ghez C, Favilla M, Ghilardi MF, Gordon J, Bermejo R, Pullman S. 1997. Discrete and continuous planning of hand movements and isometric force trajectories. Experimental Brain Research 115:217–233.

Gulberti A, Arndt P, Colonius H. 2014. Stopping eyes and hands: evidence for non-independence of stop and go processes and for a separation of central and peripheral inhibition. Frontiers in human neuroscience 8:61.

Haggard P. 2008. Human volition: towards a neuroscience of will. Nature Reviews Neuroscience 9:934–946.

Haith AM, Bestmann S. 2020. Preparation of movement In: Poeppel D, Mangun G, Gazzaniga M, editors. The Cognitive Neurosciences. MIT Press.

Haith AM, Pakpoor J, Krakauer JW. 2016. Independence of movement preparation and movement initiation. Journal of Neuroscience 36:3007–3015.

Hanes DP, Carpenter RH. 1999. Countermanding saccades in humans. Vision research 39:2777–2791.

Hannah R, Aron AR. 2021. Towards real-world generalizability of a circuit for action-stopping. Nature Reviews Neuroscience 22:538–552.

He JL, Hirst RJ, Puri R, Coxon J, Byblow W, Hinder M, Skippen P, Matzke D, Heathcote A, Wadsley CG. 2021. OSARI, an Open-Source Anticipated Response Inhibition Task. Behavior research methods 1–11.

Jagadisan UK, Gandhi NJ. 2017. Removal of inhibition uncovers latent movement potential during preparation. Elife 6:e29648.

Jahanshahi M, Obeso I, Rothwell JC, Obeso JA. 2015. A fronto–striato–subthalamic–pallidal network for goal-directed and habitual inhibition. Nature Reviews Neuroscience 16:719–732.

Kaufman MT, Seely JS, Sussillo D, Ryu SI, Shenoy KV, Churchland MM. 2016. The largest response component in the motor cortex reflects movement timing but not movement type. ENeuro 3.

Kornblum S. 1973. Simple reaction time as a race between signal detection and time estimation: A paradigm and model. Perception & Psychophysics 13:108–112.

Lakens D. 2017. Equivalence tests: A practical primer for t tests, correlations, and meta-analyses. Social psychological and personality science 8:355–362.

Lappin JS, Eriksen CW. 1966. Use of a delayed signal to stop a visual reaction-time response. Journal of Experimental Psychology 72:805.

Lara AH, Elsayed GF, Zimnik AJ, Cunningham JP, Churchland MM. 2018. Conservation of preparatory neural events in monkey motor cortex regardless of how movement is initiated. Elife 7:e31826.

Leotti LA, Wager TD. 2010. Motivational influences on response inhibition measures. Journal of Experimental Psychology: Human Perception and Performance 36:430–447.

Leunissen I, Zandbelt BB, Potocanac Z, Swinnen SP, Coxon JP. 2017. Reliable estimation of inhibitory efficiency: to anticipate, choose or simply react? European Journal of Neuroscience 45:1512–1523.

Logan GD, Cowan WB. 1984. On the ability to inhibit thought and action: A theory of an act of control. Psychological review 91:295–327.

Logan GD, Van Zandt T, Verbruggen F, Wagenmakers E-J. 2014. On the ability to inhibit thought and action: general and special theories of an act of control. Psychological review 121:66.

Logan GD, Yamaguchi M, Schall JD, Palmeri TJ. 2015. Inhibitory control in mind and brain 2.0: blocked-input models of saccadic countermanding. Psychological review 122:115.

Luce RD. 1991. Response times: Their role in inferring elementary mental organization. Oxford University Press, USA.

Matzke D, Love J, Heathcote A. 2017. A Bayesian approach for estimating the probability of trigger failures in the stop-signal paradigm. Behavior research methods 49:267–281.

Matzke D, Strickland LJG, Sripada C, Weigard AS, Puri R, He J, Hirst R, Heathcote A. 2021. Stopping timed actions. PsyArXiv. doi:https://doi.org/10.31234/osf.io/9h3v7

Matzke D, Verbruggen F, Logan G. 2018. The stop-signal paradigm. Stevens’ handbook of experimental psychology and cognitive neuroscience 5:383–427.

Mostofsky SH, Simmonds DJ. 2008. Response inhibition and response selection: two sides of the same coin. Journal of cognitive neuroscience 20:751–761.

Ollman RT, Billington MJ. 1972. The deadline model for simple reaction times. Cognitive Psychology 3:311–336.

Oosterlaan J, Logan GD, Sergeant JA. 1998. Response inhibition in AD/HD, CD, comorbid AD/HD+ CD, anxious, and control children: A meta-analysis of studies with the stop task. The Journal of Child Psychology and Psychiatry and Allied Disciplines 39:411–425.

Özyurt J, Colonius H, Arndt PA. 2003. Countermanding saccades: evidence against independent processing of go and stop signals. Perception & Psychophysics 65:420–428.

Raud L, Huster RJ, Ivry RB, Labruna L, Messel MS, Greenhouse I. 2020. A single mechanism for global and selective response inhibition under the influence of motor preparation. Journal of Neuroscience 40:7921–7935.

Rawji V, Modi S, Rocchi L, Jahanshahi M, Rothwel J. 2022. Proactive inhibition is marked by differences in the pattern of motor cortex activity during movement preparation and execution. Journal of Neurophysiology 819–828.

Salinas E, Stanford TR. 2013. The countermanding task revisited: fast stimulus detection is a key determinant of psychophysical performance. Journal of Neuroscience 33:5668–5685.

Slater-Hammel AT. 1960. Reliability, accuracy, and refractoriness of a transit reaction. Research Quarterly American Association for Health, Physical Education and Recreation 31:217–228.

Sternberg S. 1969. Memory-scanning: Mental processes revealed by reaction-time experiments. Am Sci 57:421–457.

Verbruggen F, Aron AR, Band GP, Beste C, Bissett PG, Brockett AT, Brown JW, Chamberlain SR, Chambers CD, Colonius H. 2019. A consensus guide to capturing the ability to inhibit actions and impulsive behaviors in the stop-signal task. Elife 8:e46323.

Verbruggen F, Logan G. 2017. Control in response inhibitionThe Wiley Handbook of Cognitive Control. John Wiley & Sons, Ltd. pp. 97–110.

Verbruggen F, Logan GD. 2008. Response inhibition in the stop-signal paradigm. Trends in cognitive sciences 12:418–424.

Wadsley CG, Cirillo J, Nieuwenhuys A, Byblow WD. 2022. Stopping interference in response inhibition: behavioral and neural signatures of selective stopping. Journal of Neuroscience 42:156–165.

Walker E, Nowacki AS. 2011. Understanding equivalence and noninferiority testing. Journal of general internal medicine 26:192–196.

Welford A. 1980. Choice reaction time: Basic concepts. Reaction times 73–128.

Wessel JR, Aron AR. 2017. On the globality of motor suppression: unexpected events and their influence on behavior and cognition. Neuron 93:259–280.

Wiecki TV, Frank MJ. 2013. A computational model of inhibitory control in frontal cortex and basal ganglia. Psychological review 120:329–355.

Xu J, Westrick Z, Ivry RB. 2015. Selective inhibition of a multicomponent response can be achieved without cost. Journal of Neurophysiology 113:455–465.

